# A new Karyopherin-β2 binding PY-NLS epitope of HNRNPH2 is linked to neurodevelopmental disorders

**DOI:** 10.1101/2023.01.20.524964

**Authors:** Abner Gonzalez, Hong Joo Kim, Brian D. Freibaum, Ho Yee Joyce Fung, Chad A. Brautigam, J. Paul Taylor, Yuh Min Chook

## Abstract

The normally nuclear HNRNPH2 is mutated in *HNRNPH2*-related X-linked neurodevelopmental disorder causing the protein to accumulate in the cytoplasm. Interactions of HNRNPH2 with its importin Karyopherin–β2 (Transportin-1) had not been studied. We present a structure that shows Karyopherin-β2 binding HNRNPH2 residues 204-215, a proline-tyrosine nuclear localization signal or PY-NLS that contains a typical R-X_2-4_-P-Y motif, ^206^RPGPY^210^, followed a new Karyopherin-β2 binding epitope at ^211^DRP^213^ that make many interactions with Karyopherin-β2 W373. Mutations at each of these sites decrease Karyopherin-β2 binding affinities by 70-100 fold, explaining aberrant accumulation in cells and emphasizing the role of nuclear import defects in the disease. Sequence/structure analysis suggests that the new epitope C-terminal of the PY-motif, which binds Karyopherin-β2 W373, is rare and thus far limited to close paralogs HNRNPH2, HNRNPH1 and HNRNPF. Karyopherin-β2 W373, a HNRNPH2-binding hotspot, corresponds to W370 of close paralog Transportin-2, a site of pathological variants in patients with neurodevelopmental abnormalities, suggesting that Transportin-2-HNRNPH2/H1/F interactions may be compromised in the abnormalities.

**Summary:** HNRNPH2 variants in *HNRNPH2*-related X-linked neurodevelopmental disorder aberrantly accumulate in the cytoplasm. A structure of Karyopherin-β2•HNRNPH2 explains nuclear import defects of the variants, reveals a new NLS epitope that suggests mechanistic changes in pathological variants of Karyopherin-β2 paralog Transportin-2.

## Introduction

Karyopherin-β2 and Karyopherin-β2b (Kapβ2 and Kapβ2b, also known as Transportin-1/TNPO1 or Transportin-2/TNPO2) are 85% identical in sequence and both proteins are members of the Karyopherin-β family of nuclear transport receptors (Gorlich and Kutay, 1999; Guttinger et al., 2004; Tran et al., 2007; Weis, 2003; Wing et al., 2022). Kapβ2 and Kapβ2b transport many of the same RNA binding proteins from the cytoplasm into the nucleus. These cargoes include HNRNPs A1, A2, D, F, H1, H2, FUS, EWS and TAF15, many of which are linked to neurodegenerative, neuromuscular or neurodevelopmental diseases (Chook and Suel, 2011; Suel et al., 2008). FUS, EWS, TAF15, HNRNP A1 and HNRNP A2 are linked to amyotrophic lateral sclerosis (ALS) and frontal temporal dementia (FTD), HNRNPs A1 and A2 are also involved in multisystem proteinopathy, HNRNPDL in limb girdle muscular dystrophy, and HNRNPs H1 and H2 (also known as H’) in neurodevelopmental disorders (Dormann et al., 2010; Shorter, 2019). Pathogenesis of these diseases involves aberrant cytoplasmic localization of nuclear import cargoes due to defects in Kapβ2-mediated nuclear import (Dormann et al., 2010; Zhang and Chook, 2012).

The Kapβ2 proteins bind to very diverse 15-100 residues long nuclear localization signals (NLSs) in their cargoes that are named the proline-tyrosine- or PY-NLSs. These signals reside in intrinsically disordered regions of the cargos, have overall basic character, and contain a set of 2-3 Kapβ2-binding sequence motifs or epitopes (Cansizoglu et al., 2007; Lee et al., 2006; Suel et al., 2008). PY-NLS epitope 1 is a hydrophobic or basic motif at the N-terminus of the NLS, epitope 2 is often a single arginine residue or sometimes a helix with multiple arginine residues and epitope 3 is most often a proline-tyrosine (PY) dipeptide. Epitopes 2 and 3 together make up the C-terminal RX_2-5_PY motif. Some PY-NLSs contain and use only a subset of the three epitopes to bind Kapβ2/Kapβ2b (Soniat and Chook, 2016; Suel et al., 2008). Many familial ALS and neurodevelopmental mutations in the cargoes FUS, HNRNPH2 and HNRNPH1 are found in their PY-NLSs and affect epitopes 2 or 3 (Bain et al., 2016; Bain et al., 2021; Dormann et al., 2010; Pilch et al., 2018; Reichert et al., 2020; Zhang and Chook, 2012).

Disease mutations in the FUS PY-NLS have been examined structurally or quantitatively but the mechanism of how the PY-NLS of HNRNPH2 binds Kapβ2 and how pathogenic variants affect the interaction have not been studied (Gonzalez et al., 2021; Zhang and Chook, 2012). HNRNPH2 and its close paralog HNRNPH1 are RNA processing proteins that shuttle between the nucleus and cytoplasm, but the proteins mostly reside in the nucleus (Van Dusen et al., 2010). Both proteins are involved in transcription, mRNA splicing, translation, mRNA degradation and localization. Most patients with *HNRNPH2*-related X-linked neurodevelopmental disorder have mutations in the PY-NLS of the HNRNPH2 protein (R206W, R206Q, R206G, R206L, P209L, Y210C, R212S, R212T, R212G and P213L) that cause the normally nuclear HNRNPH2 to accumulate in the cytoplasm and associate with stress granules upon stress (Fig. 1 A) (Bain et al., 2016; Bain et al., 2021; Korff et al., 2022). The mutation sites at R206 and P209 are likely in epitopes 2 and 3 of the PY-NLS, respectively, but the roles of the nearby R212 and P213 are unknown. Mutations in the PY-NLS of the closely related HNRNPH1 have also been found in patients with *HNRNPH1*-related syndromic intellectual disability (Pilch et al., 2018; Reichert et al., 2020). Here, we examined how the HNRNPH2 PY-NLS binds Kapβ2, how binding energy is distributed across the NLS sequence and how disease-causing mutations affect Kapβ2-cargo interactions.

**Figure 1.**
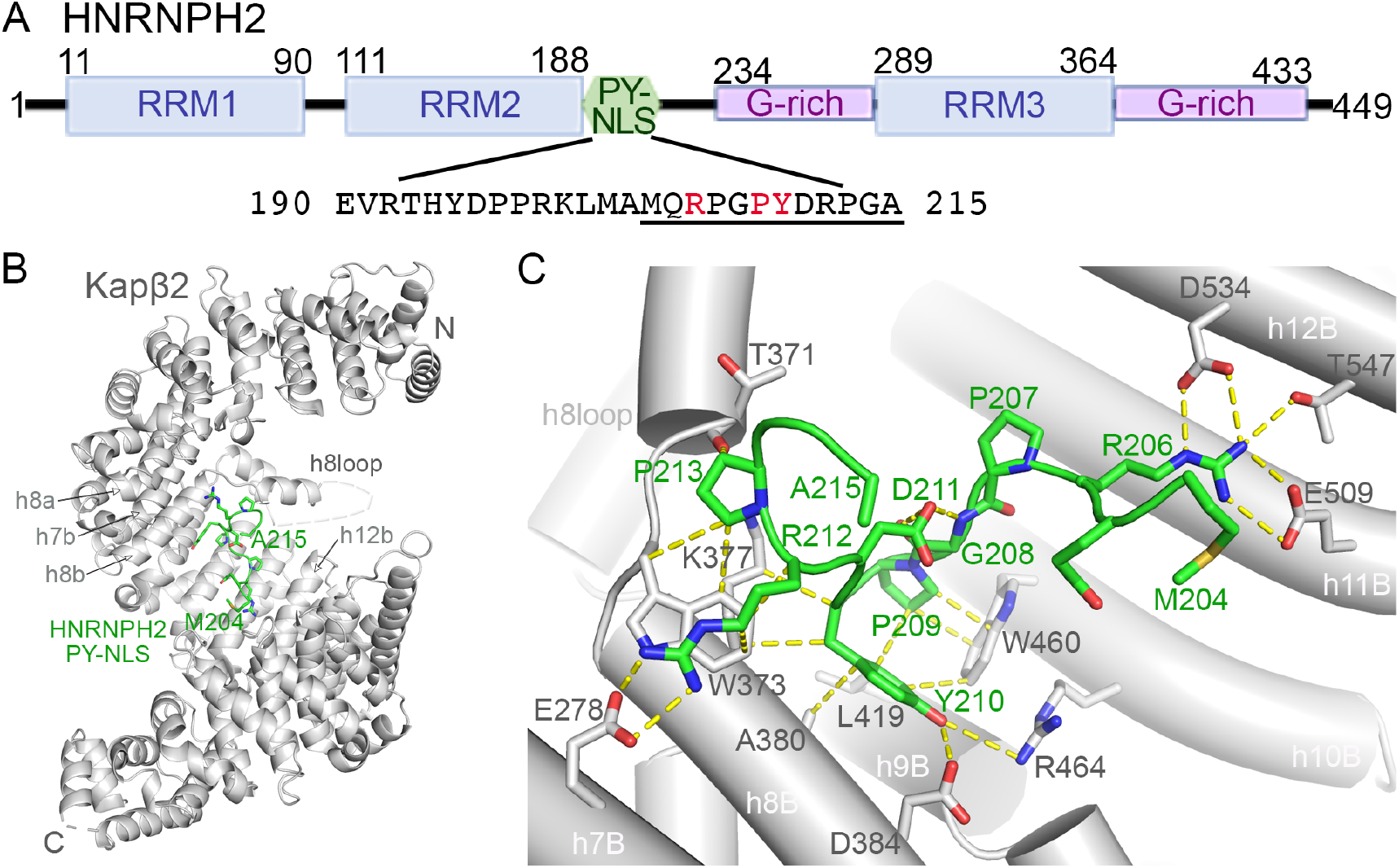
Structure of Kapβ2 bound to the HNRNPH2 PY-NLS. **(A)** Schematic of the HNRNPH2 domains and the PY-NLS sequence. PY-NLS residues that were modeled in the Kapβ2•HNRNPH2 cryo-EM structure are underlined. **(B)** Overall structure of HNRNPH2(103-225) (green) bound to Kapβ2 (gray). **(C)** Details of the HNRNPH2 PY-NLS (green) bound to Kapβ2 (gray); interactions (4Å or less) between residues are shown with yellow dashed lines.

## Results and discussion

### Cryo-EM structure of Kapβ2 bound to HNRNPH2(103-225)

The 449-amino acid HNRNPH2 protein contains three RNA recognition motif (RRM) domains: RRM1 and RRM2 are connected by a short linker while RRM2 and RRM3 are connected by a 100-reside disordered linker that contains a PY-NLS followed by a glycine-rich segment (Fig. 1 A). Using pull-down binding assays, we mapped the Kapβ2-binding HNRNPH2 segment that is most stable against proteolytic degradation to residues 103-225, which covers the RRM2 domain followed by 47 residues that contain the PY-NLS (Fig. S 1, A-C). HNRNPH2(103-225) shows minimal proteolytic degradation and binds Kapβ2 tightly with a dissociation constant (K_D_) of 50 nM (measured by isothermal titration calorimetry or ITC; Table 1 and Fig. S1 D). However, the RRM2 domain alone shows no detectable Kapβ2-binding and although the PY-NLS alone (residues 190-225) is highly prone to degradation, it also binds Kapβ2 tightly with a K_D_ of 40 nM (Table 1 and Fig. S1, E and F). Inclusion of the RRM2 in the HNRNPH2(103-225) construct appears critical for the technical purpose of maintaining an intact and non-degraded PY-NLS.

**Table 1.** ITC analysis of Kapβ2 binding to MBP-HNRNPH2.

| **Kapβ2** | **MBP-HNRNPH2** | **K<sub>D</sub> [2σ]<sup>a</sup>** |
| --- | --- | --- |
| WT | RRM2-PY-NLS (103-225) WT | 50 [24, 87] nM |
| WT | PY-NLS (190-225) WT | 40 [24, 60] nM |
| WT | RRM2 (103-189) WT | No binding |
| WT | RRM2-PY-NLS(103-225) R206W | 5.2 [2, 28] μM |
| WT | RRM2-PY-NLS(103-225) R206Q | 3.6 [1.6, 9] μM |
| WT | RRM2-PY-NLS(103-225) P209L | 16.3 [2.3, 47] μM |
| WT | RRM2-PY-NLS(103-225) R212A | 3.2 [2, 5.5] μM |
| W373A | RRM2-PY-NLS(103-225) WT | 6.1 [3, 36] μM |

<sup>a</sup>95% confidence interval determined by error-surface projection in the global analysis of duplicate experiments.

Initial attempts of cryo-EM determination with full length Kapβ2 bound to HNRNPH2(103-225), yielded a map with no clear density for the HNRNPH2 peptide. Thus, we subjected the complex to mild crosslinking and was able to solve its cryo-EM structure to 3.2 Å resolution (Table S1 and Fig. S2 A). Kapβ2 adopts a superhelical conformation of 20 HEAT repeats (h1-h20, each with a pair of antiparallel A and B helices) (Fig. 1 B). Cryo-EM density is present to model HNRNPH2 residues 204-215, which bind across the B helices of repeats h7-h12 in the C-terminal concave surface of Kapβ2 (Fig. 1 B, C and S2 B). The conformation of this binding site is very similar to those of other PY-NLS bound Kapβ2 structures (2H4M, 2OT8, 2Z5K, 4FDD, 4JLQ, 4OO6; root mean square deviation or RMSD values aligning residues in h7-h16 are 0.9-1.5 Å) (Cansizoglu et al., 2007; Imasaki et al., 2007; Lee et al., 2006; Soniat et al., 2013; Zhang and Chook, 2012). No density is present for the RRM2 domain (residues 103-188) and residues 189-203 at the N-terminus of the PY-NLS.

Many Kapβ2•PY-NLS structures in the Protein Data Bank (PDB) were solved using a Kapβ2 construct where the long and flexible 67-residue HEAT repeat 8 loop (h8loop) was truncated (Cansizoglu et al., 2007; Lee et al., 2006; Soniat and Chook, 2016; Soniat et al., 2013; Zhang and Chook, 2012). Our structure is of a complex that contains the full length Kapβ2 with an intact h8loop. We modeled Kapβ2 h8loop residues 308-325 and 362-374 that contain two short helices proximal to helices A and B of repeat h8. No density is present for distal h8loop residues 326-361 (Fig. S2 C).

### Kapβ2-HNRNPH2 PY-NLS interactions: epitopes 2 and 3

The persistently bound portion of the HNRNPH2 PY-NLS (residues 204-215) covers PY-NLS epitope 2 (R206), epitope 3 (^209^PY^210^) and five residues C-terminal of the PY-motif (^211^DRPGA^215^) (Fig. 1 C, S2 B). The absence of cryo-EM density for residues N-terminal of M204 suggests that epitope 1 of the HNRNPH2 PY-NLS either binds very weakly or is absent. HNRNPH2 PY-NLS epitope 2 or R206 is a mutational hot spot for *HNRNPH2*-related X-linked neurodevelopmental disorder (Bain et al., 2016; Bain et al., 2021). Like all previously observed epitope 2 arginine residues, R206 makes multiple salt-bridges to several acidic residues of Kapβ2 (Cansizoglu et al., 2007; Huber and Hoelz, 2017; Imasaki et al., 2007; Lee et al., 2006). ITC data show that two of the most prevalent mutations of R206 found in patients, R206W and R206Q, decreased Kapβ2 affinities by 70-100 fold (Table 1, Fig. S3, A and B). Such large Kapβ2-binding defects explain the aberrant accumulation of HNRNPH2 R206 variants in the cytoplasm and association with stress granules upon stress (Korff et al., 2022). The less prevalent R206G and R206L epitope 2 variants also cannot participate in electrostatic interactions with Kapβ2 and are expected to have substantially decreased Kapβ2 affinities and aberrant subcellular localization.

Residues that flank HNRNPH2 R206 (M204, Q205, P207 and G208) make no contact with Kapβ2, but the HNRNPH2 ^209^PY^210^ dipeptide is a typical PY-NLS epitope 3 that makes numerous hydrophobic interactions with Kapβ2 residues W460, L419 and A380 (Fig. 1 C). The side chain hydroxyl of HNRNPH2 Y210 also makes hydrogen bond-like polar interactions with Kapβ2 (Fig. 1 C). Variants of the PY motif, P209L and Y210C, have been identified in patients (Bain et al., 2016; Bain et al., 2021). The P209L mutation of HNRNPH2(103-225) decreased Kapβ2 affinity by ∼200 fold (Table 1, Fig. S3 C), consistent with its aberrant localization in cells (Korff et al., 2022). Since Y210 also makes many contacts with Kapβ2, the HNRNPH2 Y210C variant most likely also decreases Kapβ2 affinity substantially.

### Kapβ2-HNRNPH2 PY-NLS interactions C-terminal of the PY-motif

C-terminal of the PY motif, the HNRNPH2 polypeptide chain turns and almost folds back on itself with the side chain carboxylate of D211 making intramolecular hydrogen bond-like interactions with the main chain amide and carbonyl groups of G208. These intramolecular interactions likely stabilize the kinked PY-NLS conformation (Fig. 1, B and C). The turned HNRNPH2 chain positions the next two residues, R212 and P213, to make many interactions with Kapβ2 residues in repeats h7 and h8. R212 makes salt bridges with Kapβ2 residue E278 and hydrophobic interactions with Kapβ2 residue W373, while P213 interacts with Kapβ2 residues T371 and W373. The many contacts between HNRNPH2 ^212^RP^213^ and Kapβ2 stand out because Kapβ2-NLS interactions C-terminal of the PY motifs are rare. Of the structures of Kapβ2 bound to PY-NLSs of ten different cargoes in the PDB, only the Kapβ2-bound PY-NLSs of HNRNPM, NXF1 and Nab2 adopt conformations where residues C-terminal of the PY or the homologous PL motifs come close to Kapβ2 (Fig. 2, A-E). Residues C-terminal of the PY motifs of the other PY-NLSs are either flexible and not modeled, or the PY residues are the C-termini of the cargo proteins (Fig. 2, F-H) (Cansizoglu et al., 2007; Huber and Hoelz, 2017; Imasaki et al., 2007; Lee et al., 2006; Soniat et al., 2013; Zhang and Chook, 2012). The many contacts between HNRNPH2 ^212^RP^213^ and Kapβ2 are thus unusual and suggest that these two HNRNPH2 residues might be potential binding hot spots, and thus explain disease variants R212T, R212G and P213L. Indeed, mutation of R212 to alanine decreased Kapβ2 affinity by 64-fold (K_D_ of 3.2 μM; Table 1, Fig. S3D). Pull-down binding assays show that single residue variants of ^212^RP^213^ in patients, R212T, R212G and P213L, all decreased Kapβ2 binding substantially (Fig. 3 A). Kapβ2 W373, which makes many interactions with both HNRNPH2 R212 and P213, also contributes substantial binding energy as the Kapβ2(W373A) mutant binds 120-fold weaker (K_D_ 6.1 μM) (ITC data in Table 1 and Fig. S3 E). In summary, Kapβ2-HNRNPH2 interactions are dominated by strong epitopes 2 (R206), 3 (^209^PY^210^) and a new epitope 4 at ^212^RP^213^. Amino acids at these epitopes, R206, P209, Y210, R212 and P213, are sites of variants found in patients with *HNRNPH2*-related X-linked neurodevelopmental disorder. Mutations at these sites all decrease Kapβ2-HNRNPH2 interactions and Kapβ2-mediated nuclear import.

**Figure 2.**
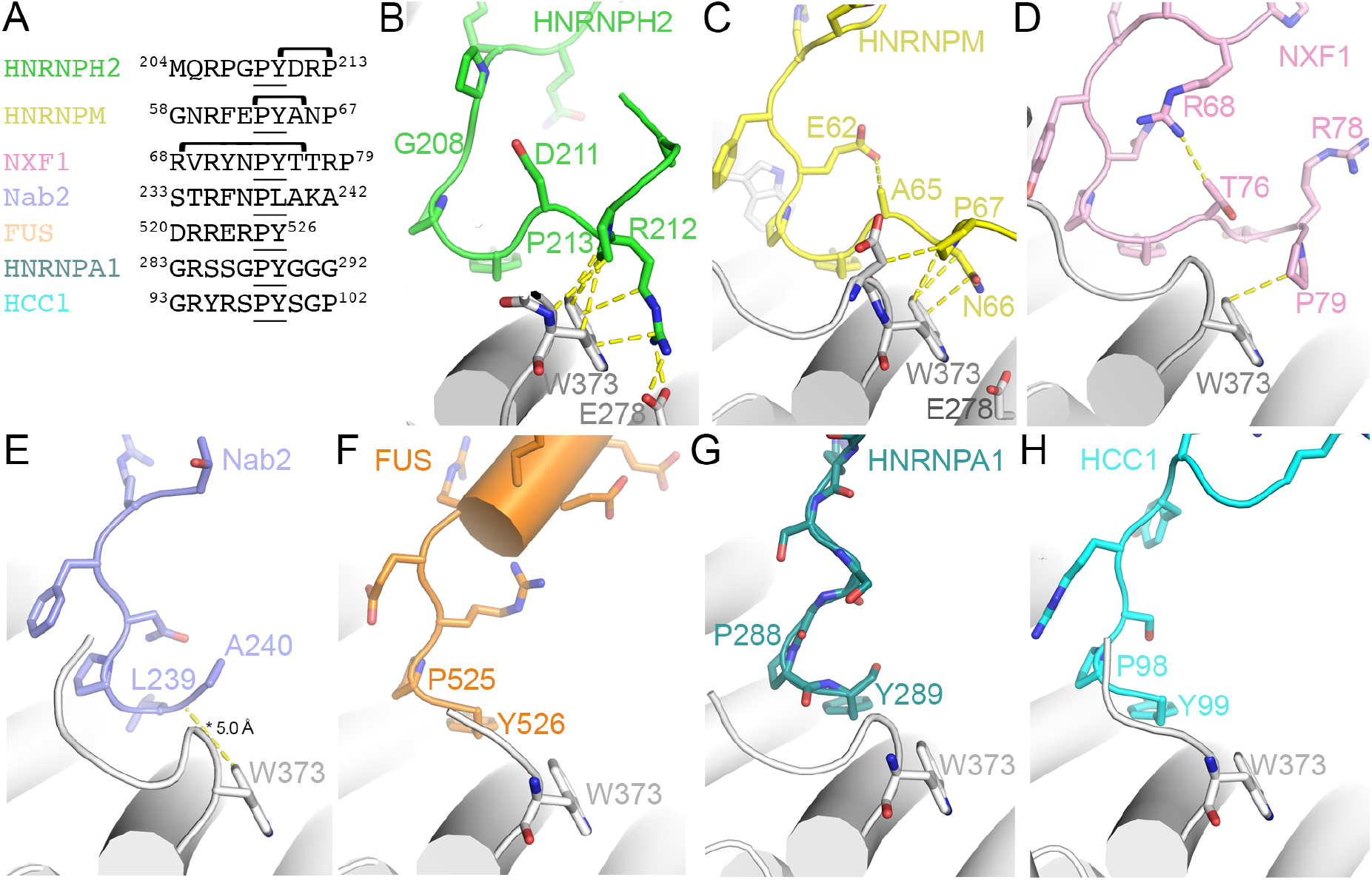
Interactions of Kapβ2 with diverse PY-NLSs. **(A)** Sequences of the C-terminal regions of PY-NLSs from HNRNPH2 (green), HNRNPM (yellow), NXF1 (pink), Nab2 (purple), FUS (orange), HNRNPA1 (teal) and HCC1 (cyan). **(B-H)** Interactions between Kapβ2 and PY-NLS residues C-terminal the PY motifs of HNRNPH2 (**B**), HNRNPM (**C**), NXF1 (**D**), Nab2 (**E**), FUS (**F**), HNRNPA1 (**G**) and HCC1 (**H**), are shown with dashed lines (interactions are < 4.0 Å except when marked with an asterisk *).

**Figure 3.**
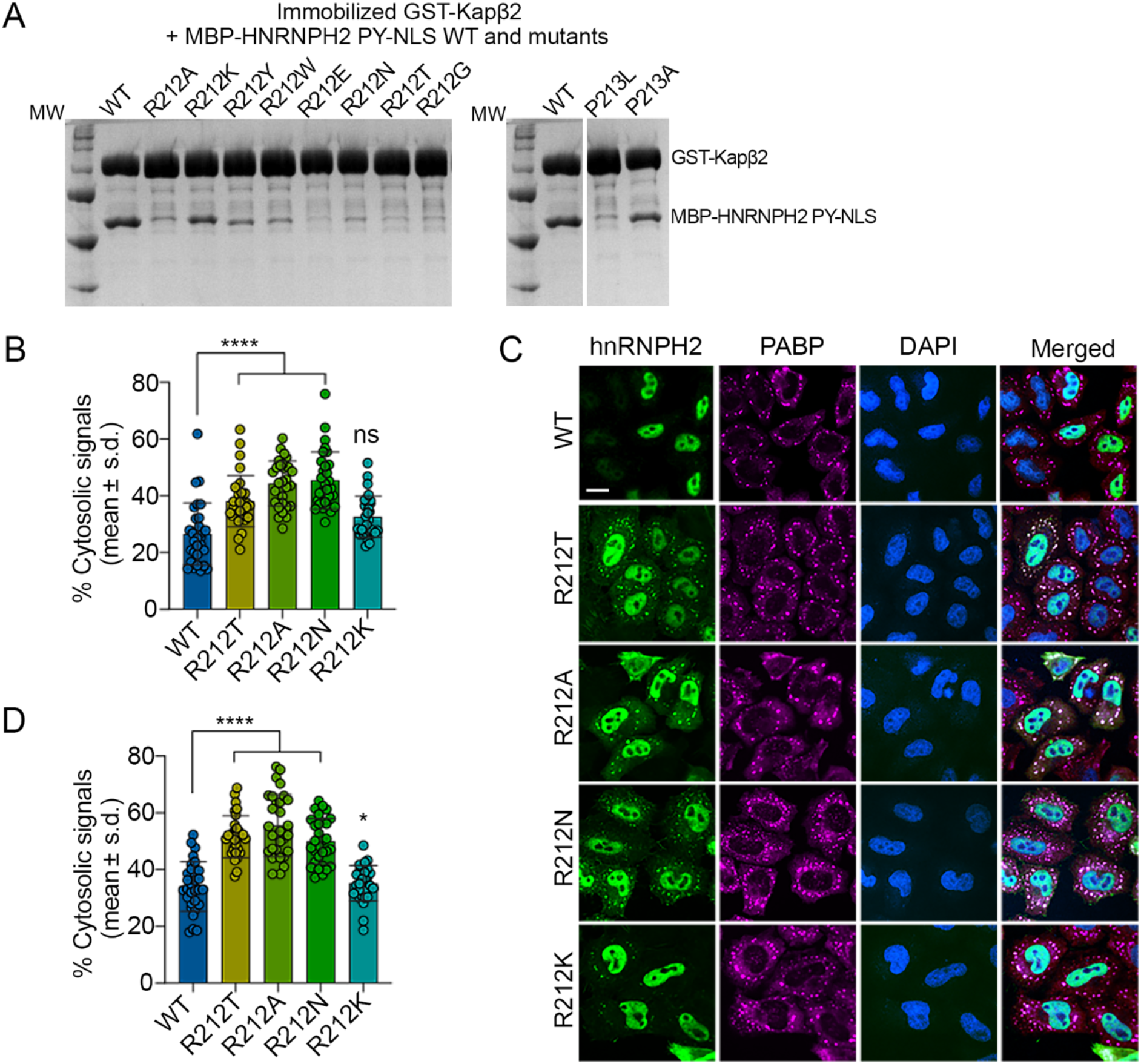
Epitope 4 of the PY-NLS affects Kapβ2-binding and nuclear transport of HNRNPH2. **(A)** Pull-down binding assays of immobilized GST-Kapβ2 binding to HNRNPH2 mutants. Following extensive washing, bound proteins are separated and visualized by SDS-PAGE and Coomassie-staining. WT control was duplicated for ease of visual comparison for the P213 mutants. **(B)** Quantification of the percentage of cytoplasmic FLAG-epitope tagged signal from cells expressing either WT hnRNPH2 or respective mutants. Forty-eight hours post-transfection, HeLa cells were fixed and stained with indicated antibodies. n=30 per condition. Error bars represent mean ± s.d., *\*\*\*\*P* < 0.0001 by one-way ANOVA with Dunnett’s multiple comparisons test. **(C-D)** Representative images (**C**) and quantification (**D**) of cytoplasmic immunofluorescent signals from cells expressing WT hnRNPH2 or respective mutants after NaAsO_2_ induction. Forty-eight hours post-transfection, HeLa cells were treated with 0.5 mM NaAsO_2_ for 30 min, fixed, and stained with indicated antibodies. PABP was used as a stress granule marker. n=30 per condition. Error bars represent mean ± s.d., *\*\*\*\*P* < 0.0001 and *\*P* = 0.0391 by one-way ANOVA with Dunnett’s multiple comparisons test. Scale bar, 10 μm.

### Epitope 4 of the PY-NLS

Residues ^66^NP^67^ of HNRNPM and residue P79 of NXF1 occupy the same positions as HNRNPH2 ^212^RP^213^ while the main chain between Nab2 residues 239-240 is positioned slightly further away from Kapβ2 (Fig. 2, A-E) (Cansizoglu et al., 2007; Imasaki et al., 2007; Soniat et al., 2013) Like HNRNPH2 ^212^RP^213^, HNRNPM ^66^NP^67^ and NXF1 P79 make multiple contacts with Kapβ2, especially with residue W373. We propose that these NLS residues constitute novel epitopes 4 of the PY-NLSs that specifically contact W373 of Kapβ2.

Like PY-NLS epitopes 1, 2 and 3, all of which can have variable contributions to total binding energies in different PY-NLSs, epitopes 4 are similarly variable (Cansizoglu et al., 2007; Lee et al., 2006; Suel et al., 2008). Many contacts of the HNRNPH2 epitope 4 are with residue W373 of Kapβ2 and almost all contacts of the HNRNPM epitope are with W373. However, while Kapβ2 W373 makes critical contributions to HNRNPH2-binding (K_D_ 6 μM for Kapβ2(W373A) binding to HNRNPH2(103-225)), Kapβ2 W373 is not important for binding the HNRNPM PY-NLS (K_D_ 9 nM for Kapβ2(W373A) binding to HNRNPM PY-NLS; Table 1, Fig. S3 F). These results suggest that epitope 4 of the HNRNPH2 PY-NLS is a Kapβ2-binding hot spot but the same epitope of HNRNPM PY-NLS is weak. Mutagenesis of NXF1 residues 76-80 (^76^TTRPN^80^) to alanines by Zhang et al. also showed that they contribute little to Kapβ2-binding, suggesting that the epitope 4 of NXF1 is also weak (Zhang et al., 2011).

We examined epitopes 4 of HNRNPH2 (^206^RPGPY<u>DRP</u>^213^; epitope 4 underlined), HNRNPM (^60^RFEPY<u>ANP</u>^67^) and NXF1 (^68^RVRYNPY<u>TTRP</u>^79^) for common features. All three sequences share a proline 2-3 residues C-terminal of the PY-motif, and residues between this C-terminal proline and the PY-motif make multiple interactions with Kapβ2. Immediately C-terminal of the PY-motif, residue D211 of HNRNPH2 may contribute to the strong epitope 4 in this PY-NLS by making intramolecular hydrogen-bond like interactions with the main chain at G208 that may stabilize or pre-organize the β-turn-like conformation at the C-terminus of the NLS (Fig. 2 B). The equivalent residue in NXF1, T76, also likely makes intramolecular hydrogen-bond-like interactions with the R68 side chain, and residue A65 of HNRNPM makes intramolecular Van der Waals contacts with E62 (Fig. 2, C and D). The next residue(s) in the epitopes 4 of HNRNPH2 (^212^RP^213^), HNRNPM (^66^NP^67^) and NXF1 (^77^TRP^79^) vary in their interactions with Kapβ2. HNRNPH2 R212 is clearly important for a strong epitope 4, as it makes electrostatic and hydrophobic interactions with Kapβ2 and mutation to alanine decreased Kapβ2 affinity substantially (Fig. 2 B, Table 1 and Fig. S3 A). HNRNPH2 P213 also makes multiple contacts with Kapβ2 W373. We mutated HNRNPH2 R212 to lysine, asparagine, threonine, tyrosine, tryptophan, glutamic acid, gycine and alanine to test the importance of both hydrophobic and electrostatic interactions in epitope 4. Binding assays show that the R212A, R212Y, R212W, R212E, R212N, R212T and R212G mutations decreased the amounts of HNRNPH2 PY-NLS pulled down by immobilized Kapβ2, consistent with decreases in Kapβ2-binding. R212K only mildly the amount of HNRNPH2 pulled down by Kapβ2 (Fig. 3 A). These data suggest that both hydrophobic and electrostatic interactions from basic residues are important for Kapβ2-binding at epitope 4, explaining why basic side chains like arginine, lysine and likely histidine at this position are important for a strong epitope 4. We also mutated HNRNPH2 P213 to leucine as this is a disease variant (Fig. 3 A). The P213L mutation abolished HNRNPH2 pull-down by Kapβ2, consistent with decreased Kapβ2-binding, explaining aberrant accumulation in cells and a role for nuclear import defects in the disease (Korff et al., 2022). Interestingly, a P213A mutation decreased Kapβ2-binding only slightly, suggesting that although a proline at this position is ideal, a small side chain like alanine can be tolerated.

To validate the consequence of epitope 4 mutations on subcellular localization of HNRNPH2, we expressed FLAG-epitope tagged hnRNPH2 WT and R212 variants (R212A, R212N, and R212K) in addition to the disease variant R212T in HeLa cells. As previously observed (Korff et al., 2022), the R212T variant had increased cytoplasmic accumulation at basal condition (Fig. 3 B) and further accumulation in the cytoplasm and stress granules after exposure to 30 minutes of sodium arsenite (Fig. 3, C and D). The R212A and R212N mutants which were found to have impaired Kapβ2-binding *in vitro* have similar cytoplasmic accumulation to the disease mutant at both basal and stress conditions (Fig. 3, A-D). The R212K variant, which only mildly decreased Kapβ2-binding *in vitro* was found to have similar levels of cytoplasmic hnRNPH2 protein with wild type protein at both conditions (Fig. 3, A-D), suggesting that binding Kapβ2 through R212 residue is critical for nuclear import of HNRNPH2.

The residues equivalent of HNRNPH2 ^212^RP^213^ in NXF1 (^77^TRP^79^) do not contact Kapβ2. Here, although T76 is likely engaged in a hydrogen-bond like intramolecular interaction, the non-optimal orientation of ^77^TRP^79^ for contacts with Kapβ2 W373 may explain this weak epitope 4 (Zhang et al., 2011). Having two residues between the PY and the C-terminal proline appears suitable for positioning the side chain that precedes the latter to contact Kapβ2 W373. Of the equivalent ^66^NP^67^ of HNRNPM, residue N66 makes hydrophobic but not electrostatic or polar interactions with Kapβ2 (Fig. 2, C and D). The non-ideal intramolecular interaction of the preceding side chain (A65) and the few intermolecular interactions of HNRNPM ^66^NP^67^ are thus insufficient to make the PY-NLS epitope 4 a strong epitope (Table 1, Fig. S3 F) (Cansizoglu et al., 2007).

Based on the above results and sequence/structural analysis, we propose a consensus sequence ξ-R/K/H-P/A (ξ, hydrophilic amino acid) that should immediately follow the PY-motif to predict a strong epitope 4 of the PY-NLS.

### Epitope prevalence, Kapβ2 tryptophan residues in disease and PY-NLS specificity

PY-NLS sequences of ten different cargos for which Kapβ2-bound structures are available are shown in Fig. 4 A along with sequences of close paralogs. We also examined the sequences of other PY-NLSs that were reported to bind Kapβ2 (Fig. 4 B) for the predicted strong epitope 4 motif (β-R/K/H-P) or for a looser likely weak motif (X_1-2_-R/K/H-P where X is a non-polar amino acid) that immediately follows the PY-motif. No “strong” epitope 4 motif is present in the sequences in Fig. 4 B and potentially “weak” epitope 4 motifs are present in only two PY-NLSs. This finding suggests that strong epitopes 4 are uncommon in PY-NLSs. Of the 40 PY-NLS sequences in Fig. 4, A and B, the only strong epitopes 4 identified or predicted are in HNRNPH2 and close paralogs HNRNPH1 and HNRNPF (Fig. 4 A). If PY-NLS epitope 4 is uncommonly used in Kapβ2 cargos, we expect that its binding site on the karyopherin (W373) is also not often used for cargo-binding.

**Figure 4.**
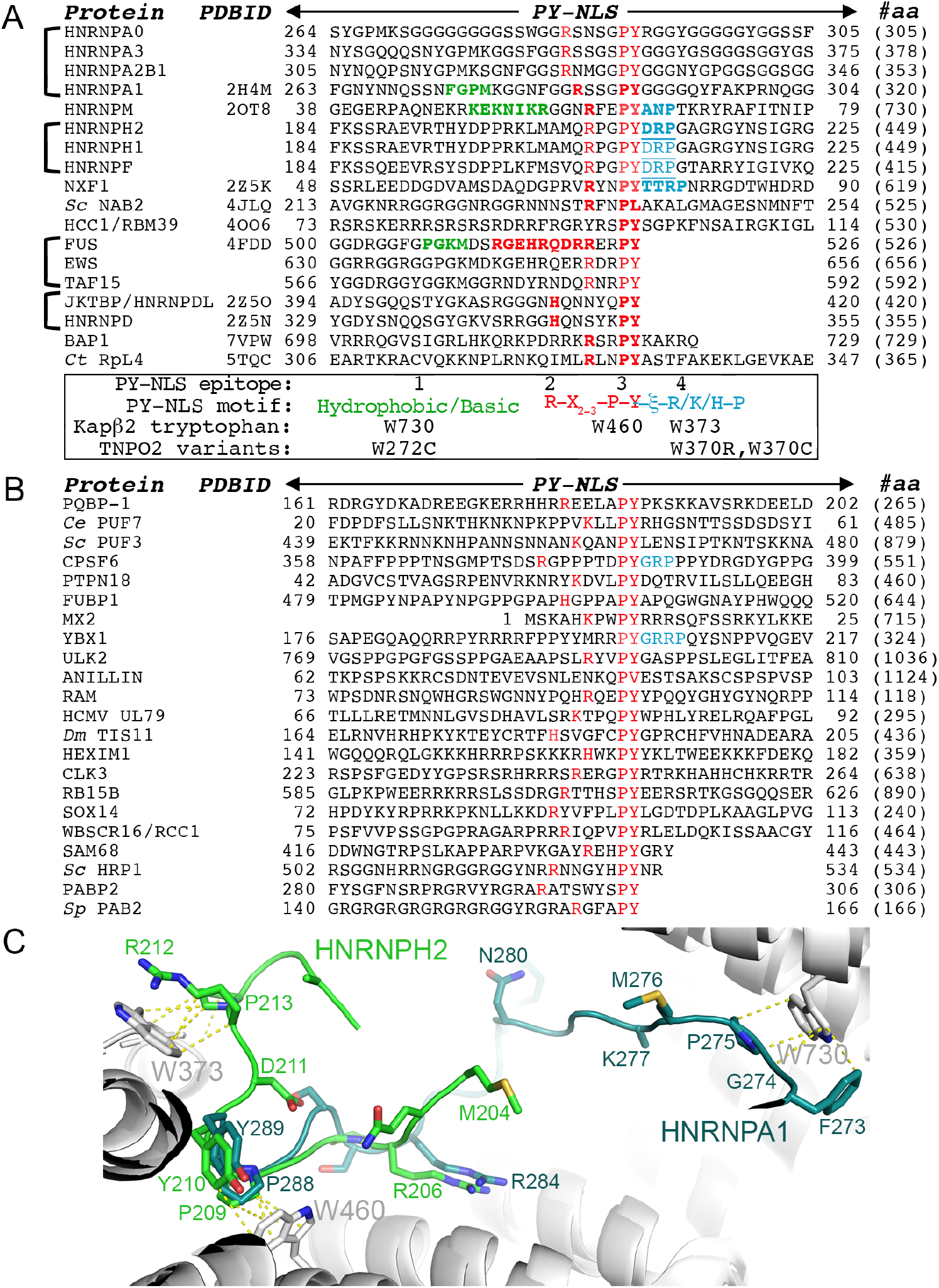
PY-NLS sequences and their Kapβ2-binding epitopes. **(A)** The sequences of PY-NLSs with structures bound to Kapβ2 and the sequences of their close paralogs. The residues of epitopes 1 that are observed in the structures are colored green and residues of the R/K/H-X_2-5_-P-Y/L motifs (epitopes 2 and 3) are colored red. Observed/predicted epitopes 4 are colored blue. **(B)** Previously reported PY-NLS sequences with residues of the R/K/H-X_2-5_-P-Y/L motifs colored red and the predicted epitopes 4 colored blue. **(C)** Structures of Kapβ2 (gray) bound to hnRNPH2 (green) and HNRNPA1 (teal) (Kapβ2 of the twp structures aligned). Tryptophan residues that interact with the PY-NLS epitope 1 (W730), epitope 3 (W460) and epitope 4 (W373) are shown. Interactions with the tryptophan residues are shown with dashed lines.

Examination of Kapβ2-PY-NLS structures and the distributions of binding energies across the PY-NLSs suggests that epitope 1 (N-terminal basic/hydrophobic motif) is also rarely strong (Cansizoglu et al., 2007; Lee et al., 2006; Suel et al., 2008; Zhang and Chook, 2012). Epitope 1 is not modeled in many structures and of the ones resolved structurally, the epitope is strong only in HNRNPA1 but weak in HNRNPM and in FUS (Cansizoglu et al., 2007; Zhang and Chook, 2012). In contrast, epitopes 2 and 3, which make up the C-terminal R-X_2-3_-P-Y/ϕ motif (ϕ is hydrophobic), are often used and the PY dipeptides often contribute substantially to Kapβ2-binding (Cansizoglu et al., 2007; Suel et al., 2008; Zhang and Chook, 2012).

We note that epitopes 1 of PY-NLSs contact W730 on Kapβ2 helix h16B, epitopes 3 contact W460 on helix h10B and epitopes 4 contact W373 which is immediately N-terminal of helix h8B (Fig. 4 C). The WW/AA mutant of Kapβ2, in which W460 and W730 are mutated to alanines, is commonly used to test cargo-binding (Barraud et al., 2014; Lee et al., 2006). Our assessment that epitopes 1 and 4 are used sparsely in Kapβ2 cargos is interesting given the recent identification of variants at the epitopes’ binding sites on Kapβ2b/TNPO2 found in patients with neurodevelopmental abnormalities (Goodman et al., 2021). W370 of TNPO2 (homologous to Kapβ2 W373; epitope 4) is mutated to arginine or cysteine and W727 (homologous to Kapβ2 W730; epitope 1) mutated to cysteine (Goodman et al., 2021). These TNPO2 variants will likely be defective in importing only a small subset of PY-NLS containing cargos including HNRNPH2, HNRNPH1 and HNRNPF, and nuclear import defects that result from TNPO2 W370R and W370C in neurodevelopmental abnormalities patients may be related to those in *HNRNPH2*-related X-linked neurodevelopmental disorders.

## Conclusion

The structure of the HNRNPH2 PY-NLS bound to Kapβ2 show an NLS that spans residues 204-215. Epitope 1 or the N-terminal hydrophobic/basic motif of the PY-NLS is either missing or weak and therefore not persistently bound to Kapβ2. Instead, the PY-NLS of HNRNPH2 consists of energetically strong epitopes 2 (R206), 3 (^209^PY^210^) and a newly defined epitope 4 (^212^RP^213^). Mutations at each of these epitopes, corresponding to different pathogenic variants, decrease binding affinities for Kapβ2 by 70-100 fold, explaining their mislocalization to the cytoplasm of cells. Strong epitopes 4 are not common in Kapβ2 cargos; they are so far found only in HNRNPH2 and its close paralogs HNRNPH1 and HNRNPF. Epitope 4 makes many interactions with Kapβ2 W373, corresponding to pathological variants at W370 of the close paralog TNPO2 that cause neurodevelopmental abnormalities.

## Methods

### Protein expression and purification

The plasmid to overexpress GST-Kapβ2 was as described in previous work (Chook and Blobel, 1999). Various truncation variants of HNRNPH2 described in this work were subcloned into pMal-TEV with or without His_6_ inserted before the MBP, or into the pGex-tev vector as for the GST-Kapβ2 overexpression plasmid. Some single amino acid mutants were generated by site-directed mutagenesis and others were purchased synthesized (GenScript).

GST-Kapβ2 was overexpressed in BL21-Gold (DE3) *E. coli* cells (#230132, Agilent Technologies) and expression was induced with 0.5 mM isopropyl-β-d-1-thiogalactoside (IPTG; #I2481C, Goldbio) for 12 h at 25 °C. The bacterial cells were harvested by centrifugation, resuspended in lysis buffer (50 mM Tris pH 7.4 (#T60040, RPI), 150 mM NaCl (#S23020, RPI), 1 mM EDTA (#E57020, RPI), 2 mM β-mercaptoethanol (#M6250, Sigma-Aldrich), 15% v/v glycerol (#G7893, Sigma-Aldrich), 1 mM benzamidine (#434760, Sigma-Aldrich), 10 μg/mL leupeptin (#J61188, Alfa Aesar) and 50 μg/mL AEBSF (#A540, Goldbio) and then lysed with the EmulsiFlex-C5 cell homogenizer (Avestin, Ottawa, Canada). GST-Kapβ2 proteins were purified by affinity chromatography using Glutathione Sepharose 4B beads (GSH; #17075604, Cytiva). For ITC analysis and cryo-EM structure determination, the GST tag was removed by adding TEV protease to GST-Kapβ2 on the GSH column. Kapβ2 released from the GSH beads was further purified by anion exchange chromatography followed by size-exclusion chromatography (Superdex 200 increase, Cytiva). For pull-down binding assays, GST-Kapβ2 was eluted from GSH beads with 20mM glutathione at pH 6.5 (#G4251, Sigma-Aldrich) and the protein was further purified by anion exchange followed by size-exclusion chromatography (Superdex 200 Increase).

MBP-HNRNPH2 and His_6_-MBP-HNRNPH2 proteins were overexpressed in BL21-Gold (DE3) *E. coli* cells (induced with 0.5 mM IPTG for 16 h at 20 °C). The bacteria cells were lysed in buffer containing 50 mM HEPES pH 7.4 (#H400, Goldbio), 1.5 M NaCl, 10% v/v glycerol, 2 mM β-mercaptoethanol and cOmplete protease inhibitor cocktail (#05056489001, Sigma-Aldrich). The high salt is used to disrupt association with nucleic acids. MBP-HNRNPH2 proteins were purified by affinity chromatography using amylose resin (#E8021, New England BioLabs) and eluted with buffer containing 20 mM HEPES pH 7.4, 150 mM NaCl, 2 mM DTT, 10% v/v glycerol, and 20 mM maltose (#M5885, Sigma-Aldrich). His_6_-MBP-HNRNPH2 proteins were first purified using Ni-NTA Agarose (#30230, Qiagen) and eluted with buffer containing 250 mM imidazole pH 7.8 (#79227, Sigma-Aldrich), 10 mM NaCl, 10% v/v glycerol, 2mM β-mercaptoethanol. Both MBP-HNRNPH2 and His_6_-MBP-HNRNPH2 proteins were further purified by size-exclusion chromatography (Superdex 200 Increase).

### Pull-down binding assays for Kapβ2 binding to immobilized GST-HNRNPH2 proteins

*E. coli* (BL21-Gold) transformed with pGEX-tev plasmids expressing GST-HNRNPH2 proteins were grown to OD_600_ 0.6. Protein expression was then induced with 0.5 mM IPTG for 4 hours at 37°C. Cells were harvested by centrifugation, resuspended in lysis buffer (50 mM Tris pH 7.4, 150 mM NaCl, 1 mM EDTA, 2 mM DTT, 15% glycerol, 1 mM benzamidine, 10 μg/mL leupeptin and 50 μg/mL AEBSF) and lysed by sonication. The lysate was centrifuged and the supernatant containing GST-HNRNPH2 proteins added to Glutathione Sepharose 4B beads. The solution was spun down to obtain bead bed with immobilized GST-HNRNPH2 proteins, which was then washed with 1ml of lysis buffer. 50 μl of bead slurry containing ∼60 μg immobilized GST-HNRNPH2 proteins were then incubated with 8 μM Kapβ2 in 100 μl total volume for 30 min at 4°C and then washed three times with 1 ml of lysis buffer. Proteins bound on the beads were eluted by boiling in SDS sample buffer and visualized by Coomassie staining of SDS-PAGE gels.

### Isothermal titration calorimetry

Kapβ2 and MBP-HNRNPH2 proteins were dialyzed into ITC buffer (20 mM Tris pH 7.5, 150 mM NaCl, 10% v/v glycerol, 2 mM β-mercaptoethanol). ITC experiments were performed in a MicroCal PEAQ-ITC (Malvern Panalytical, Worcestershire, UK) calorimeter; it has a stirred 206.2 μL reaction cell held at 20 ºC. The first injections were 0.5 μL, followed by twenty 1.9 μL injections with a stirring rate of 750 rpm. Kapβ2 was used at 10 μM in the ITC cell; 100-350 μM MBP-HNRNPH2 proteins were used in the syringe. All ITC experiments were performed in duplicate. ITC data were integrated and baseline corrected using NITPIC (Keller et al., 2012). The integrated data were globally analyzed in SEDPHAT (Houtman et al., 2007) using a model considering a single class of binding sites. Thermogram and binding figures were plotted in GUSSI (Brautigam, 2015).

### Cryo-EM sample and grid preparation

Kapβ2 and HNRNPH2(103-225) were mixed at room temperature at 1:1.4 molar ratio, followed by a rapid addition of glutaraldehyde (#16100, Electron Microscopy Sciences) to a final concentration of 0.025% for 1 minute, and immediate injection to size-exclusion chromatography in a Superdex 200 Increase column. Fractions containing the crosslinked mixture were pooled, aliquoted and stored at −80°C for later use. Aliquots were diluted to a approximate concentration of 2.7 mg/mL in buffer containing 20 mM Tris-HCl pH 7.5, 150 nM NaCl, 2 mM β-mercaptoethanol and 0.003125% [w/v] NP-40 (#S226, Biovision Detergents Pak I) to set up cryo-EM grids. 4 μL of Kapβ2•HNRNPH2 was applied to a 300 mesh copper grid (Quantifoil R1.2/1.3) that was glow-discharged using a PELCO easiGlow glow discharge apparatus at 30 mA/30 s on top of a metal grid holder (Ted Pella). Excess sample was blotted 3 s before plunge-freezing in a Vitrobot System (Thermo Fisher) at 4°C with 95% humidity.

### Cryo-EM data collection and data processing

Cryo-EM data collection for the Kapβ2•HNRNPH2(103-225) complex was collected at the UT Southwestern Cryo-Electron Microscopy Facility on a Titan Krios at 300 kV with a Gatan K3 detector in correlated double sampling super-resolution mode at a magnification of 105,000x corresponding to a pixel size of 0.415 Å. Each movie was recorded for a total of 60 frames over 5.4 s with an exposure rate of 8 electrons/pixel/s. The datasets were collected using SerialEM (Mastronarde, 2005) software with a defocus range of −0.9 and −2.4 μm.

A total of 5,937 movies were collected for Kapβ2•HNRNPH2. The dataset was processed using cryoSPARC (Punjani et al., 2017) where it was first subjected to Patch Motion Correction (F-crop factor= 1/2) and Patch CTF Estimation. The Blob Picker was implemented on 25 micrographs to pick all possible particles with little bias. This small set of particles were subjected to 2D Classification to generate 2D templates. A subset of templates was selected and used in Template Picker, resulting in 4,042,358 particles selected. 681,236 particles of Kapβ2•HNRNPH2 were selected after 14 rounds of 2D Classification and were then sorted into seven 3D classes using Ab-initio reconstruction followed by Heterogeneous Refinement. The 208,572 particles from one 3D class of the Kapβ2•HNRNPH2 complex were utilized for Non-uniform Refinement which yielded a 3.17 Å resolution map.

### Cryo-EM model building, refinement, and analysis

The Kapβ2 and HNRNPH2 proteins were built using coordinates from the deposited structure PDB:2OT8. Model was roughly docked into the map using UCSF Chimera (Pettersen et al., 2004) and then subjected to real-space refinement with global minimization and rigid body restraints on Phenix (Adams et al., 2010). The resulting models were then manually rebuilt and refined using Coot (Emsley and Cowtan, 2004), further corrected using ISOLDE (Croll, 2018) on UCSF ChimeraX (Goddard et al., 2018), and subjected to more rounds of refinement in Phenix. UCSF ChimeraX and PyMOL version 2.5 were used for 3D structure analysis (Schrödinger, http://www.pymol.org/pymol).

### Cellular localization analysis of HNRNPH2 proteins

HeLa cells were grown in Dulbecco’s modified Eagle’s medium (DMEM) supplemented with 10% fetal bovine serum (FBS). Cells were counted using ADAM-CellT (#ADAM-CellT, NanoEntek Inc., Seoul, Korea), plated and transfected using ViaFect (#E4981, Promega) for transient overexpression according to the manufacturer’s instructions.

HeLa cells were seeded on 4-well glass slides (Millipore). Forty-eight hours post transfection for overexpression, cells were stressed with 500 μM sodium arsenite (#S7400, Sigma-Aldrich) for 30 min. Cells were then fixed with 4% paraformaldehyde (#15710, Electron Microscopy Sciences), permeabilized with 0.5% Triton X-100 (#22140, Electron Microscopy Sciences), and blocked in 3% bovine serum albumin (# A8806, Sigma-Aldrich). Primary antibodies used were mouse monoclonal anti-FLAG (1:1000, M2; #F1804, Sigma-Aldrich) and rabbit polyclonal anti-PABP (1:1000; #ab21060, Abcam). For visualization, the appropriate host-specific Alexa Fluor 488 or 647 secondary antibodies (#A21202 and A31573, Invitrogen) were used. Slides were mounted using Prolong Gold Antifade Reagent with DAPI (#P36931, Invitrogen). Imaging was performed using a Yokogawa CSU W1 spinning disk attached to a Nikon Ti2 eclipse with a Photometrics Prime 95B camera using Nikon Elements software (version 5.21.02). The DAPI and PABP channels were used to segment the nucleus and cytoplasm using the freehand selection tool on ImageJ. Integrated intensity of the nucleus, and integrated cellular signal was quantified and background signal subtracted. Integrated cytoplasmic signal was calculated by subtracting the integrated nuclear signal from the integrated cell signal. Percent cytoplasmic signal was calculated by dividing the integrated cytoplasmic signal over the integrated cell signal.

## Supplemental material

Table S 1 contain cryoEM data and model statistics. Fig. S 1 contain data for the formation of Kapβ2•HNRNPH2 complex. Fig. S 2 shows various views of the map and model with local resolution and FSC curve. Fig. S 3 contain additional ITC data.

## Acknowledgements

We thank the Structural Biology Laboratory and the Cryo-EM Facility at UTSW, which are partially supported by grant RP170644 from the Cancer Prevention & Research Institute of Texas (CPRIT), for their assistance with cryo-EM data collection. We thank the Macromolecular Biophysics Resource at UTSW for training and use of their ITC resources. This work was funded by NIGMS of NIH under Awards R35GM141461 (Y.M.C.), R01GM069909 (Y.M.C.), the Welch Foundation Grants I-1532 (Y.M.C.), support from the Alfred and Mabel Gilman Chair in Molecular Pharmacology and the Eugene McDermott Scholar in Biomedical Research (Y.M.C.), the Howard Hughes Medical Institute (J.P.T) and by NS of NIH under awards R35NS097974 (J.P.T) and F31NS120465 (A.G.).

## Author contributions

Conceptualization, Y.M.C.; Methodology and data validation, Y.M.C., A.G., H.J.K., B.F., C.A.B., J.P.T.; Investigation, A.G., B.F.; Writing, Y.M.C, A.G., H.Y.J.F, B.F., H.J.K.

## Supplementary Materials

**Table S1:** Cryo-EM data collection, refinement and validation.

| | | Kapβ2-HNRNPH2(103-225) |
| --- | --- | --- |
| **Data collection and processing** |  |  |
| Facility |  | UTSW |
| Magnification |  | 105kx |
| Voltage (kV) |  | 300 |
| Electron exposure (e<sup>-</sup>/Å<sup>2</sup>) |  | 60 |
| Defocus range (μm) |  | -1.0 to -2.2 |
| Pixel size (Å) |  | 0.83 |
| Symmetry imposed |  | C1 |
| Initial particle images (no.) |  | 4,042,358 |
| Final particle images (no.) |  | 208,572 |
| Map resolution (Å) |  | 3.17 |
| FSC threshold |  | 0.143 |
| **Refinement** |  |  |
| Initial model used (PDB code) |  | 2OT8 |
| Model resolution (Å) |  | 3.04/3.12/3.34 |
| FSC threshold |  | 0/0.143/0.5 |
| Map sharpening *B* factor (Å<sup>2</sup>) |  | 116.5 |
| Model composition |  |  |
| Nonhydrogen atom |  | 6858 |
| Protein residues |  | 860 |
| Ligand |  | 0 |
| *B* factors (Å<sup>2</sup>) |  |  |
| Protein |  | 45.24 |
| Ligand |  | 0 |
| R.m.s. deviations |  |  |
| Bond lengths (Å) |  | 0.003 |
| Bond angles (°) |  | 0.604 |
| **Validation** |  |  |
| MolProbity score |  | 1.21 |
| Clashscore |  | 4.3 |
| Poor rotamers (%) |  | 0 |
| Ramachandran plot |  |  |
| Favored (%) |  | 99.06 |
| Allowed (%) |  | 0.94 |
| Outliers (%) |  | 0.0 |

**Figure S1.**
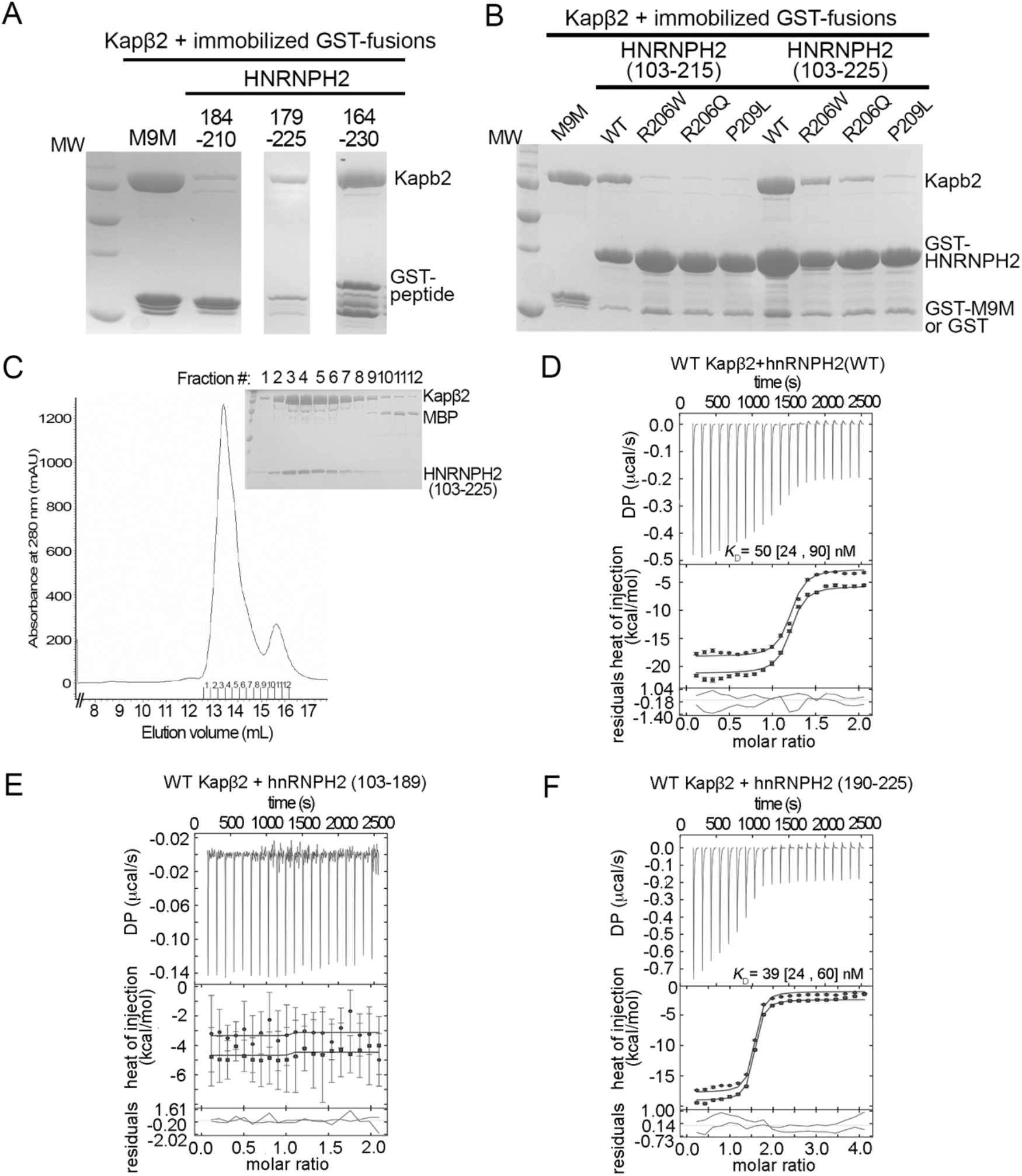
Assembly of the Kapβ2-HNRNPH2 complex. **A)** Pull-down binding assay using immobilized GST-HNRNPH2 constructs of different lengths against Kapβ2. **B)** Pull-down binding assay using immobilized GST-HNRNPH2(103-215) and GST-HNRNPH2(103-225) against Kapβ2. **C)** Size-exclusion chromatograph and SDS-PAGE gel of collected fractions showing Kapβ2 complexing with HNRNPH2(103-225). **D-F)** ITC analysis of Kapβ2 binding to MBP fusions of HNRNPH2 fragments. GUSSI output image for the global analysis of Kapβ2 binding to MBP-HNRNPH2 constructs. Top panel illustrate the reconstructed thermogram from NITPIC, middle panel shows the binding isotherm and bottom panel shows the residuals. All experiments were performed in duplicate on PEAQ-ITC. Dissociation constants were displayed with 95% confidence intervals in brackets.

**Figure S2.**
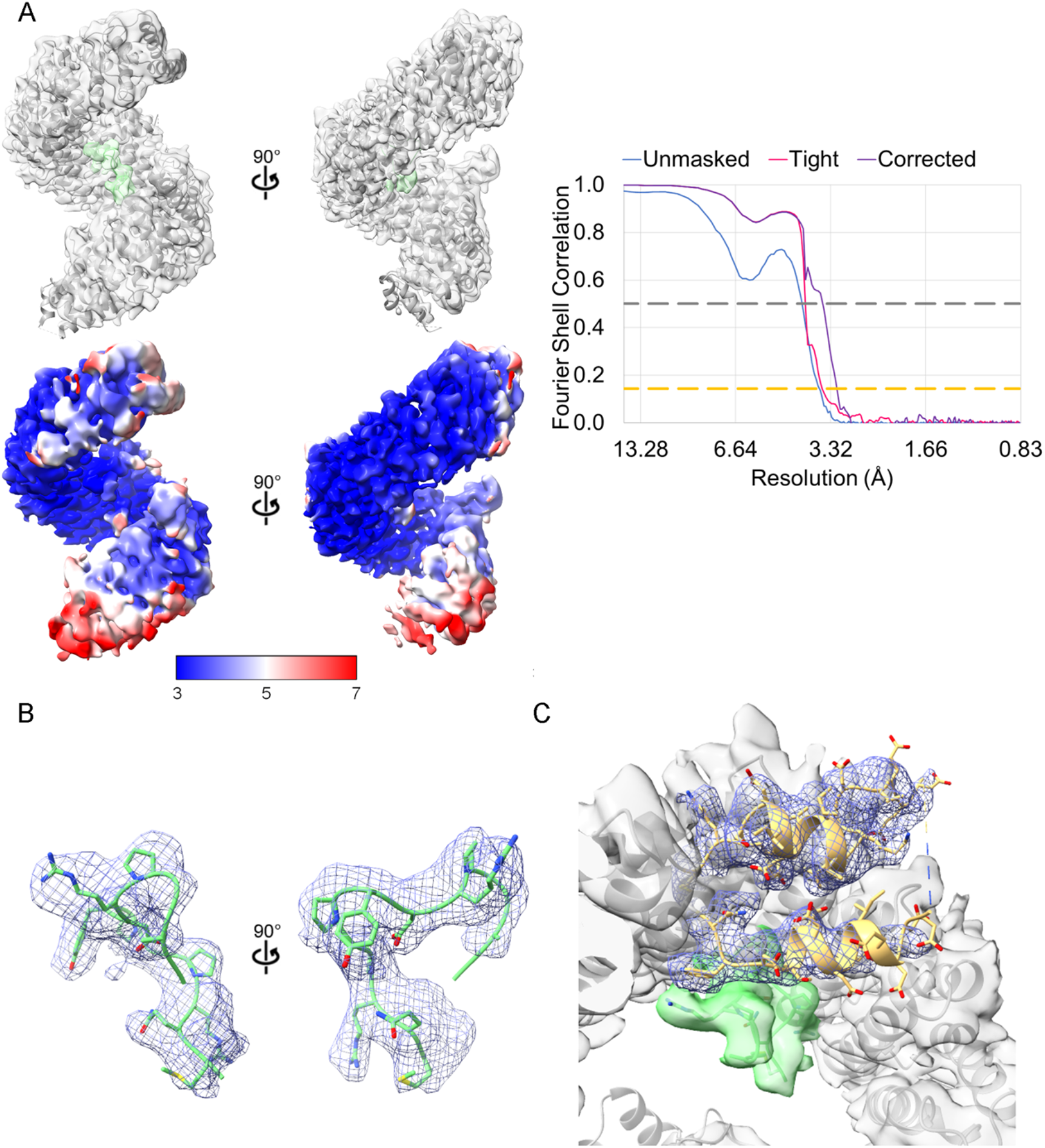
Cryo-EM structure of Kapβ2•HNRNPH2(103-225). **A)** Top left, the Kapβ2•HNRNPH2 model docked into its map with Kapβ2 in gray and the HNRNPH2 PY-NLS in green. Bottom left, the local resolution map of the complex colored accordingly to the scale. To the right, the FSC curve for the Kapβ2•HNRNPH2 PY-NLS complex cryo-EM map (unmasked in blue, tight in pink and corrected in purple). The gray dashed line represents the FSC=0.5 and the gold dashed line represents the FSC=0.143. **B)** Isolated map density (blue mesh) for the PY-NLS (green) in same orientation but zoomed view of **A**). **C)** Split map densities (blue mesh) for h8loop (yellow). The rest of the map is colored as green for PY-NLS and grey for Kapβ2. Density and model images obtained using UCSF ChimeraX.

**Figure S3.**
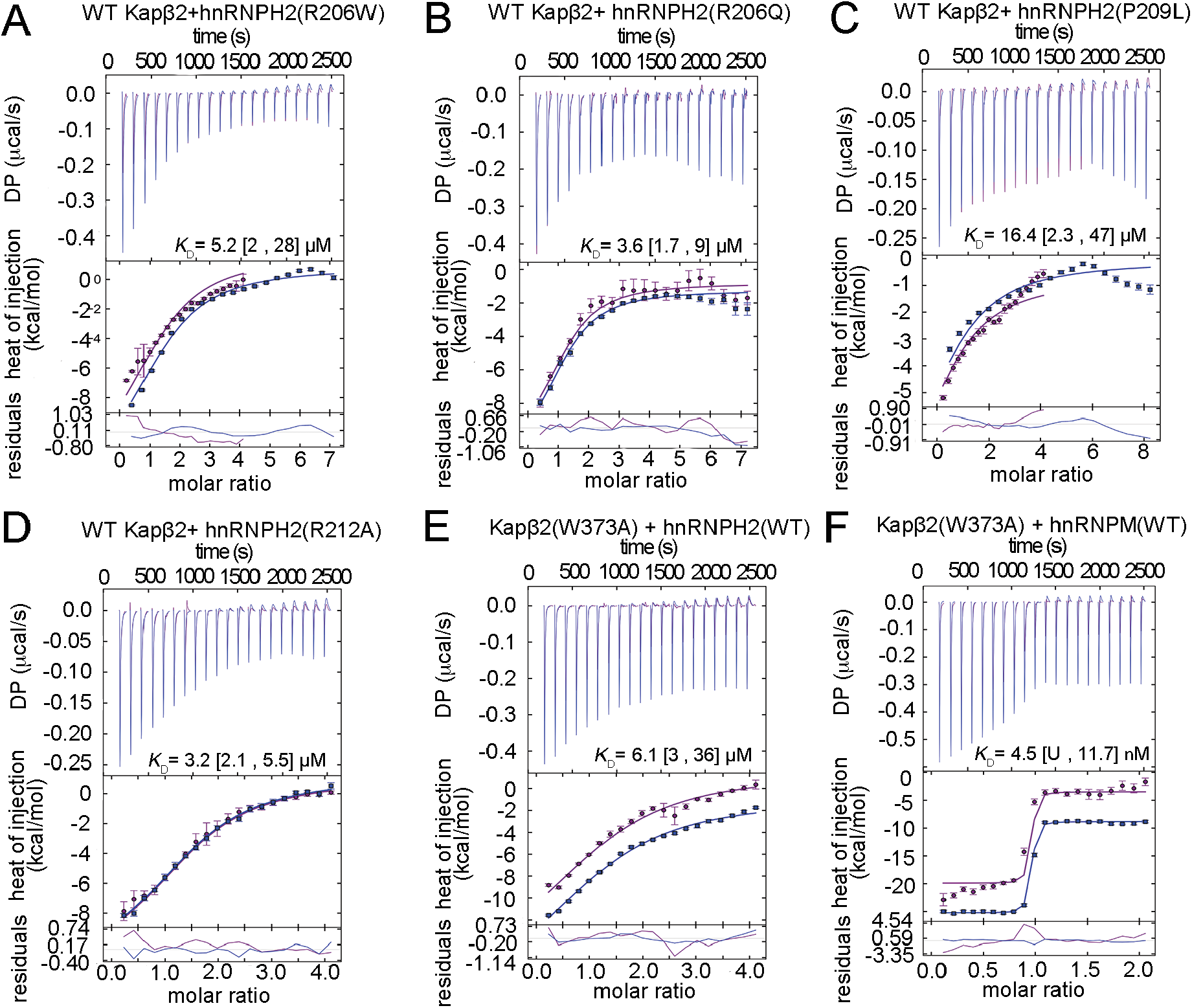
ITC analysis of Kapβ2/Kapβ2(W373A) binding to wild type or mutant HNRNPH2 and HNRNPM PY-NLSs. **A-F)** GUSSI output image for the global analysis of duplicate experiments of Kapβ2 binding to MBP-FUS constructs performed on PEAQ-ITC. Top panel illustrate the reconstructed thermograms from NITPIC, middle panel shows the binding isotherms and bottom panel shows the residuals from fitting. Dissociation constants with 95% confidence interval in brackets were displayed in the graph. U means unable to be determined.

